# Tissue-specific sex difference in the metabolism of fatty acid esters of hydroxy fatty acids

**DOI:** 10.1101/2023.11.15.567158

**Authors:** Martin Riecan, Veronika Domanska, Cristina Lupu, Maulin Patel, Michaela Vondrackova, Martin Rossmeisl, Alan Saghatelian, Florea Lupu, Ondrej Kuda

## Abstract

Fatty acid esters of hydroxy fatty acids (FAHFAs) are endogenous bioactive lipids known for their anti-inflammatory and anti-diabetic properties. Despite their therapeutic potential, little is known about the sex-specific variations in FAHFA metabolism. This study investigated the role of Androgen Dependent TFPI Regulating Protein (ADTRP), a FAHFA hydrolase. Additionally, tissue-specific differences in FAHFA levels, focusing on the perigonadal white adipose tissue (pgWAT), subcutaneous white adipose tissue (scWAT), brown adipose tissue (BAT), plasma, and liver, were evaluated using metabolomics and lipidomics. We found that female mice exhibited higher FAHFA levels in pgWAT, scWAT, and BAT compared to males. FAHFA levels were inversely related to *Adtrp* mRNA, which showed significantly lower expression in females compared with males in pgWAT and scWAT. However, no significant differences between the sexes were observed in plasma and liver FAHFA levels. *Adtrp* deletion had minimal impact on both sexes’ metabolome and lipidome of pgWAT. However, we discovered higher endogenous levels of triacylglycerol estolides containing FAHFAs, a FAHFA metabolic reservoir, in the pgWAT of female mice. These findings suggest that sex-dependent differences in FAHFA levels occur primarily in specific WAT depots and may modulate local insulin sensitivity in adipocytes. However, further investigations are warranted to fully comprehend the underlying mechanisms and implications of sex effects on FAHFA metabolism in humans.

## INTRODUCTION

Fatty acid esters of hydroxy fatty acids (FAHFAs) are a class of endogenous bioactive lipids belonging to the estolide family (LIPID MAPS [FA0709] subclass). They are defined as oligomers of a hydroxy fatty acid (HFA) and a fatty acid (FA) (1, 2). Specifically, 9-PAHSA stands for palmitic acid (PA) and hydroxy stearic acid (HSA), with the ester linkage at the “9” position relative to the carboxylic function (1). Recently, FAHFAs have attracted interest because of their remarkable anti-inflammatory, anti-diabetic, and anti-oxidant effects (1-6).

A complete understanding of the molecular mechanisms regulating FAHFAs in organisms still needs to be achieved. However, several FAHFA hydrolytic enzymes have been identified: adipose triglyceride lipase (ATGL), carboxyl ester lipase (CEL), hormone-sensitive lipase (HSL), androgen-induced protein 1 gene (AIG1), and androgen-dependent tissue factor pathway inhibitor regulating protein (ADTRP) (7-11). ATGL is a well-known lipase that can remodel FAHFAs stored in triacylglycerol estolides (TG EST) in the presence of a specific co-activator named comparative gene identification-58 (CGI-58) (11, 12) and also drives FAHFA biosynthesis (13). The putative pathway scheme (Figure 1) (14) summarizes FAHFA-related metabolic reactions and enzymes that are dependent on sex, namely ADTRP, AIG1, HSL, ATGL, and diacylglycerol O-Acyltransferase 1 (DGAT1) (15-18).

**Figure 1.**
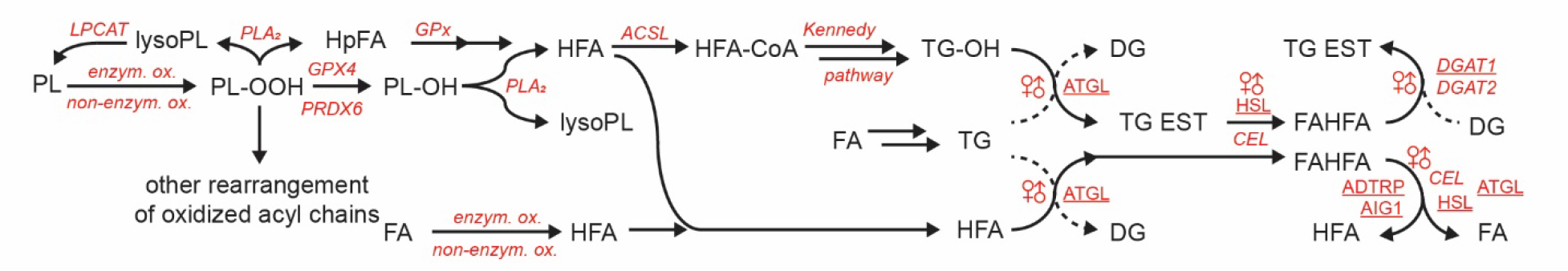
Putative scheme of FAHFA synthesis adapted from (14). Enzymes that are regulated by sex hormones are underlined. ACSL, acyl-CoA synthetase; ADTRP, Androgen-dependent TFPI-regulating protein; AIG1, Androgen-induced protein 1; ATGL, adipose triglyceride lipase; CEL, carboxyl ester lipase; DG, diacylglycerol; DGAT, diacylglycerol acyltransferase; FA, fatty acid; GPx, peroxidase activity; GPX4, glutathione peroxidase 4; HFA, hydroxy fatty acid, e.g., 13-HLA; HpFA, hydroperoxy FA; HSL, hormone-sensitive lipase; LPCAT, acyltransferase activity; lysoPL, lysophospholipid; PLA2, phospholipase A2 activity; PL, phospholipid, e.g. PC; PL-OH, hydroxylated PC; PL-OOH, peroxidated PL; TG, triacylglycerol; TG EST, triacylglycerol estolide (TG-FAHFA).

The ADTRP was first identified as a protein involved in regulating tissue factor pathway inhibitor, responsible for the anticoagulant protection of the endothelium and preservation of normal cardiovascular function (19, 20). Later studies revealed a hydrolytic activity leading to the degradation of FAHFAs (8, 9). Androgens, namely testosterone, induce transcriptional activation of ADTRP through direct binding of the androgen receptor (AR) to the androgen response element (ARE) at the ADTRP promoter region (21).

We hypothesized that since FAHFAs are degraded by enzymes whose activity is influenced by androgen levels, we should be able to find sex-dependent contrasts in FAHFA levels. We aimed to perform a lipidomic, metabolomic, and gene expression analysis in mice to define the sex-dependent metabolism of FAHFAs.

## MATERIALS AND METHODS

### Sample sources

This study used two murine models. The first animal model (MODEL 1) of transgenic *Adtrp* knockout (KO) mice was generated using CRISPR/Cas9-mediated indel mutations at the Salk Institute for Biological Studies in California as was published previously (9). Male and female WT and KO mice were maintained in a controlled environment with 12-h light/dark cycle and fed a standard chow diet with unlimited access to water in the animal facility of the Institute of Physiology, Czech Academy of Sciences. Tissue samples of perigonadal white adipose tissue (pgWAT) were collected at random fed state between 7-9 am from 4 – 6 months old mice into cryovials, rapidly frozen in liquid nitrogen, and stored at – 80 °C until extraction.

The second mouse model (MODEL 2) was bred and maintained in animal core facilities of the Oklahoma Medical Research Foundation in Oklahoma City. Mice with the null allele for the *Adtrp* gene were generated as before (22), using the bacterial artificial chromosome gene targeting strategy. We studied female and male, WT and KO, mice fed a standard chow diet. Samples of plasma, liver, pgWAT, subcutaneous WAT (scWAT), and inter scapular brown adipose tissue (BAT) were collected from ad libitum fed eight months old mice maintained in a controlled environment 12-h light/dark cycle with unlimited access to water. The mice were fasted for 5 hours starting 7 am and sacrificed by noon. Samples were frozen in liquid nitrogen and stored at – 80 °C until extraction.

### Sample extraction for lipidomic and metabolomics profiling

Tissue samples (10-25 mg) were processed using bi-phase extraction with cold methanol, methyl *tert*-butyl ether (MTBE), and water as before (14, 23). The extracts were aliquoted for the lipidomics and metabolomics profiling platforms as described in detail previously (14).

### Sample extraction for FAHFA analysis

Tissues (50-100 mg) and plasma (150 μL) samples were extracted as previously described (1, 5, 14, 23). Briefly, samples were homogenized using the bead mill in a mixture of citric acid buffer and ethyl acetate (ratio 1:2, v/v) containing one ng of [^13^C_4_]-9-PAHSA (Cayman Pharma, Neratovice, Czech Republic) as an internal standard for quantification. The organic phase was collected and dried using a Savant SpeedVac (ThermoFisher Scientific, Bremen, Germany). Extracts were resuspended in hexane and ethylacetate mixture (ratio 95:5, v/v) and purified by solid-phase extraction (SPE) using HyperSep SPE columns (500 mg/10 mL, 40–60 μm, 70 Å; Thermo Scientific). FAHFAs were eluted with ethyl acetate and dried (24). The purified extracts were resuspended in 50 μL of methanol and immediately analyzed.

### Lipidomic and metabolomic profiling

Lipidomic and metabolomic profiling of rat and mouse tissues was conducted using liquid chromatography-mass spectrometry (LC-MS) systems consisting of a Vanquish ultra-high performance liquid-chromatography (UHPLC) System (Thermo Fisher Scientific, Bremen, Germany) coupled to a Q Exactive Plus mass spectrometer (Thermo Fisher Scientific, Bremen, Germany) operated in the negative and positive ion mode, as described previously (23). Internal standards such as 1-cyclohexyluriedo-3-dodecanoic acid and Val-Tyr-Val were used to control the platform’s stability and possible differences in injection volumes (25). Obtained untargeted lipidomic and metabolomic data were processed using MS-DIAL ver. 4.92 (26). Raw data were filtered using blank samples and quality control (QC) pool samples with relative standard deviation (RSD) < 30% and then normalized using the LOESS approach utilizing QC pool samples for each matrix regularly injected between 10 actual samples (25). Lipids and metabolites were annotated using an in-house retention time–*m*/*z* library and MS/MS libraries (NIST20, LipidBlast, MassBank, MoNA). See Lipidomics Minimal Reporting Checklist (SI Material) for details.

### FAHFA regioisomer analysis

Analysis of FAHFA regioisomers was conducted on an LC-MS system consisting of a UHPLC Ultimate 3000 RSLC (Thermo Scientific) coupled with a QTRAP 5500/SelexION, a hybrid triple-quadrupole, and a linear ion trap mass spectrometer equipped with an ion mobility cell (SCIEX, Massachusetts, USA). The measurement was based on a pre-calculated multiple reaction monitoring (MRM) method using one quantifier (FA) and two qualifier transitions (HFA, HFA-H_2_O) for FAHFA (5, 27, 28). The quantifier ion was used as a survey scan for information-dependent acquisition in a linear ion trap for enhanced-resolution MS/MS and second-generation (MS/MS/MS) product ion spectra (5). This approach was used to identify FAHFA regioisomers based on the branching position of the backbone HFA using MS/MS/MS, as previously described (29). The amount of FAHFAs was quantified by normalizing its peak height to the peak height of the internal standard [^13^C_4_]-9-PAHSA obtained from MultiQuant software (SCIEX) and the total tissue weight/plasma volume (30).

### Gene expression analysis

Total RNA isolated from mouse MODEL 2 tissues using Trizol (Thermo Fisher Scientific) was purified and transcribed into cDNA as described (22). qPCR was performed using predesigned primers (Prime‐ Time, Integrated DNA Technologies Inc, Coralville, IA, USA) for mouse *Aig1* and androgen receptor (*Ar*), and RT^2^SYBR Green master mix (Qiagen, Hilden, Germany) in a CFX96 Real Time System (Bio‐Rad Labs, Inc., Hercules, CA, USA). *Adtrp* primers (forward: CCA AGA AAA TGG CCT GCA AGA G; and reverse: GAA GGA ATG AGG AGA AGC TAA GG) were designed to recognize exons 3-4, which are deleted in the MOUSE 2 model of *Adtrp*-KO (22).

### Data analysis and statistics

Data processing was performed using MetaboAnalyst 5.0 (log10 data transformation and auto-scaling) (31). GraphPad Prism 9 was used to construct bar graphs and for statistical analyses. Plotly was used to draw complex plots (32). Student’s t-test analysis and two-way ANOVA (Tukey’s multiple comparisons test) were used to compare study groups, and *p* < 0.05 was considered significant. The UpSet plot was used to visualize two-way ANOVA results (33). LORA was used to perform an over-representation analysis (34). Single-cell portal was used to explore mouse WAT data study SCP1376 (35, 36).

## RESULTS

### FAHFA levels in the pgWAT are higher in female mice

To determine whether biological sex and ADTRP affect FAHFA levels, we used targeted LC-MS methods to measure the concentration of 9-PAHSA, the most researched FAHFA representative, in pgWAT, scWAT, BAT, liver, and plasma samples (Figure 2). Consistent with previous data on MODEL 1 mice (9), the absence of ADTRP in male mice increased 9-PAHSA levels in pgWAT (Figure 2A, see Figure 2G & H for statistics). The same effect was more pronounced in female mice, which generally had higher pgWAT 9-PAHSA levels than males. Similar results were obtained in mice generated by a different knockout strategy, MODEL 2 (Figure 2B).

**Figure 2.**
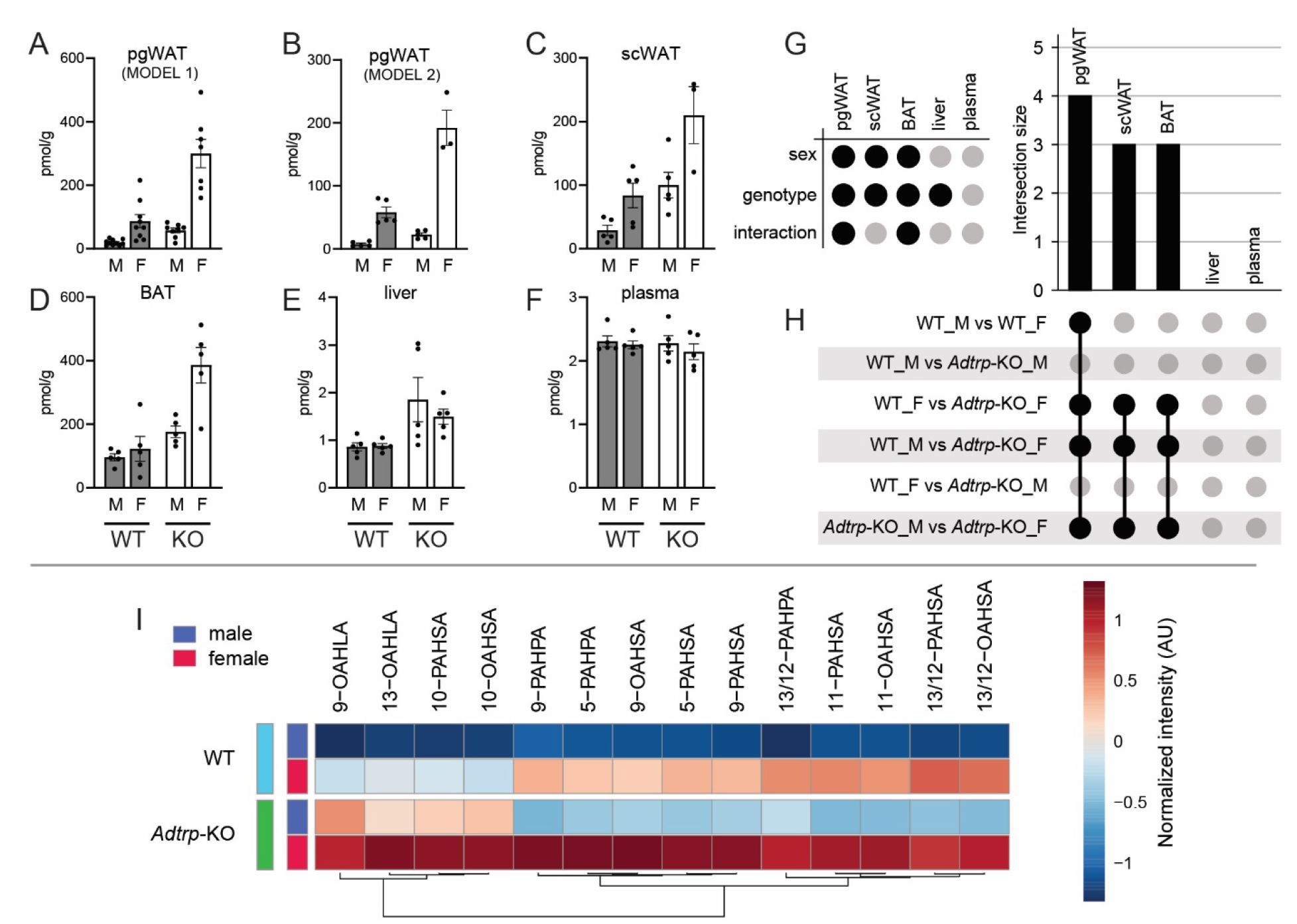
9-PAHSA levels in mouse organs. A: Levels of 9-PAHSA in pgWAT from MODEL 1 mice (9). Data are means ± SE, *n* = 7-10. Grey bars, WT mice; white bars, *Adtrp*-KO mice; M, males; F, females. B: Levels of 9-PAHSA in pgWAT from MODEL 2 mice (22). Data are means ± SE, *n* = 3-5. C-F, 9-PAHSA levels in mouse organs from MODEL 2 mice (22). Data are means ± SE, *n* = 3-5. G, Results of two-way ANOVA for panels B-F. Filled circles indicate a statistically significant effect of sex, genotype, or interaction, F-values 113.1, 52.02, 33.05 respectively, df = 1. Bonferroni’s multiple comparison test, *p* < 0.05. Panel A statistics are similar to panel B. H: UpSet plot highlighting pairwise comparison by two-way ANOVA. Filled circles connected by a line indicate statistically significantly different results. The size of the intersection indicates the number of significant tests. I. Levels of FAHFAs in pgWAT, MODEL 2. Data are means, *n* = 3-5, auto-scaled, and log10 transformed.

Next, we examined other ADTRP-expressing tissues related to lipid metabolism. Sex-related differences in 9-PAHSA levels were observed in scWAT and BAT, whereas genotype-related differences were observed in all three adipose depots and the liver (Figure 2C-E). No difference was observed in plasma (Figure 2F, fed state). Figure 2G summarizes the results of the 2-way ANOVA (sex vs. genotype), and the UpSet plot highlights the results of the multiple comparison tests. Other FAHFAs were regulated similarly to 9-PAHSA in the pgWAT (Figure 2I).

FAHFA levels were inversely related to *Adtrp* mRNA, which showed significantly lower expression in females compared with males in pgWAT and scWAT (Figure 3). *Adtrp* mRNA levels were undetectable in *Adtrp*-KO mice, but a compensatory upregulation of *Aig1* mRNA was found in *Adtrp*-KO males (Figure 3B); see also Figure 1 in (9). Androgen receptor mRNA levels were upregulated in *Adtrp*-KO animals, suggesting an imbalance in steroid signaling (Figure 3C).

**Figure 3.**
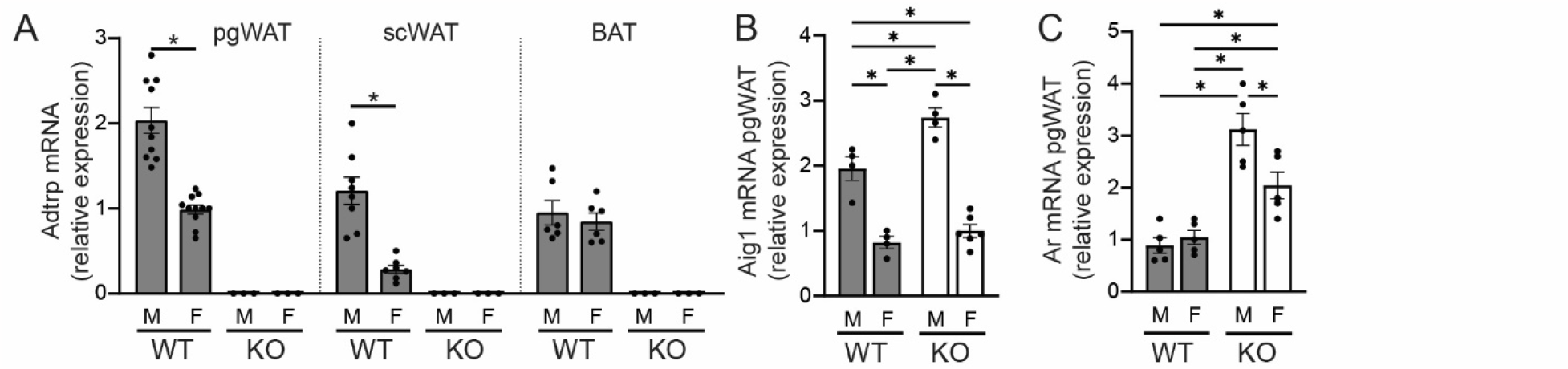
Gene expression in adipose tissue depots, MODEL 2 mice (22). A: Levels of *Adtrp* mRNA in pgWAT (t = 6.838, df = 19), scWAT (t = 5.242, df = 13) and BAT. Data are means ± SE, n = 6-11. * Statistically significant difference, *p* < 0.05, t-test. B, C: *Aig1* (F-value 121.6, 13.52, 5.323 for sex, genotype, and interaction, df = 1) and *Ar* (F-value 4.264, 51.43, 7.459 for sex, genotype, and interaction, df = 1) mRNA expression in pgWAT. Data are means ± SE, n = 4-6. * Statistically significant difference, *p* < 0.05, two-way ANOVA, Tukey’s multiple comparisons test.

### ADTRP deletion only slightly affected the metabolome and lipidome of pgWAT of both sexes

Next, we focused only on the pgWAT, where the most prominent differences between males and females were observed, and we examined the metabolome and lipidome profiles of both genotypes of MODEL 1 as the FAHFA metabolism is interlinked with phospholipid and acylglycerol metabolism (Figure 4, Table S1, Table S2). The effect of genotype on the metabolome was weak (Table S1), while the effect on the lipidome was mild (Table S2). Using loose statistical parameters (|fold change| > 1.2 and *p*-value < 0.05), we detected 41 downregulated and 1 upregulated lipid in male KO mice (Figure 4A) and 8 downregulated lipids in female KO mice (Figure 4B). The downregulated clusters consisted of glycerolipids [GL] and glycerophospholipids [GP], and the common lipids for both sexes were triacylglycerols and triacylglycerol estolides (Figure 4C). Lipid overrepresentation analysis showed that the downregulated clusters were significantly enriched with glycerolipids composed of 7 acyl chains. The lipid molecule representing the most enriched structural features was TG 18:0_18:1_20:1 (Figure 4D).

**Figure 4.**
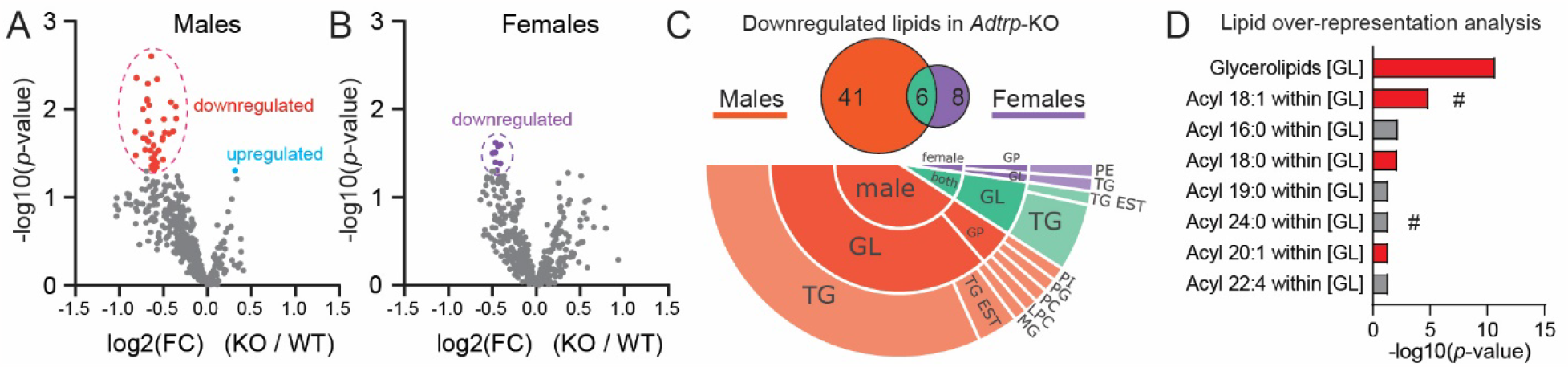
Effect of *Adtrp-*KO on the lipidome. A: Volcano plot of pgWAT lipidome from male mice, MODEL 1. The red cluster marks 41 downregulated lipids, and the blue dot marks upregulated lipids (|fold change| > 1.2 and *p*-value < 0.05, t-test). B: Volcano plot for female mice. Purple cluster marks 8 downregulated lipids. C: Venn diagram of clusters from both sexes and color-coded lipid class composition of the clusters. Abbreviations follow the LIPID MAPS nomenclature: glycerolipids [GL], and glycerophospholipids [GP] classes. D: Lipid overrepresentation analysis of downregulated clusters; # marks features specific to lipids shared by both sexes. Red bars indicate building blocks representing the most enriched structural features.

### Endogenous levels of TG estolides are higher in the pgWAT of females

The esterified form of FAHFAs, TG estolides, were only slightly affected by ADTRP deficiency in pgWAT (Figure 4), similar to what was previously reported in female mice (9). However, when we examined only the effect of sex, we found higher levels of total TG estolides in females (Figure 5A). Heatmaps of the individual TG estolide species showed clustering based on total lipid unsaturation, but the separation was not statistically significant (Figure 5B and Figure S3).

**Figure 5.**
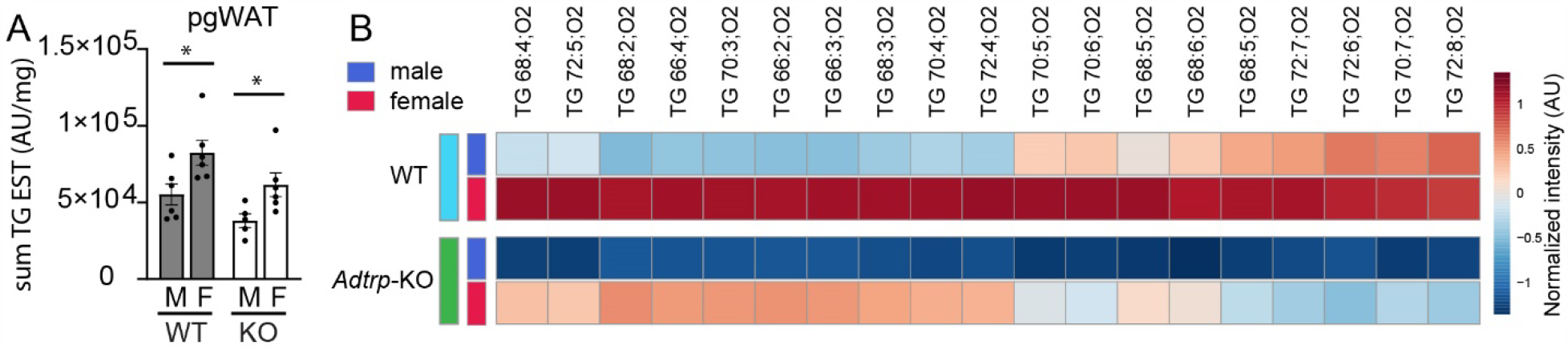
Effect of *Adtrp*-KO on TG estolides. A: The sum of TG estolide levels in pgWAT for both sexes and genotypes, MODEL 1. Data are mean ± SE, *n* = 5-6. WT, wild type; KO, *Adtrp* knockout. * Statistically significant difference between genotypes at *p* < 0.05, two-way ANOVA (F-value 12.50, 7.049, 0.07 for sex, genotype and interaction, df = 1). B: Heatmap representation of individual lipids. See Figure S1 for a detailed version. Data are auto-scaled and log10 transformed, means, *n* = 5-6.

## DISCUSSION AND CONCLUSION

Previous studies have shown that endogenous FAHFA levels in rodents and humans can be influenced by age (37), dietary composition (5, 23, 38), exercise training (30), and metabolic disorders such as obesity and diabetes (1, 39). The effect of sex on FAHFA and TG estolide levels in adipose tissue was a prominent finding in this study. The most significant effect of sex on tissue FAHFA and TG estolide levels was observed in pgWAT, probably due to its close contact with the gonads. In contrast, more distant adipose depots, such as scWAT and BAT, may be exposed to lower levels of androgens from circulation, since the local androgen-dependent *Adtrp* expression appears to be lower.

Importantly, adipose tissue has been established as an important site for steroid storage and metabolism (40). Locally produced or stored androgens could modulate the exposure of adipocytes to active androgens and potentially modulate levels of FAHFA hydrolases. The expression of steroid-converting enzymes differs in different adipose tissue depots, e.g., higher levels of 17β-hydroxysteroid dehydrogenase were found in omental than in subcutaneous adipose tissue (41). Moreover, dihydrotestosterone stimulated lipolysis and upregulated Atgl and Hsl in male rat pgWAT (18), suggesting a potential steroid effect on the FAHFA biosynthesis, degradation, and FAHFA release from TG estolides (10, 12).

The contribution of the liver to systemic FAHFA metabolism is lower than that of adipose tissue (13). We speculate that the sex-dependent FAHFA metabolism has a role at the local tissue level, e.g., modulation of local glucose uptake because the systemic 9-PAHSA levels in the circulation were not altered.

The complete absence of *Adtrp* had only a minor effect on the pgWAT metabolome and lipidome. From the global perspective, no differences between WT and *Adtrp*-KO were detected, as reported previously (9). Only the targeted approach (targeted FAHFA analysis and single metabolite analysis without the FDR correction) showed that the FAHFAs and TG estolides were altered by *Adtrp* deletion. Interestingly, the sex differences in PAHSAs persisted and were even greater upon *Adtrp* deletion, which suggests that other enzymes (see compensatory *Aig1* overexpression in KO mice) involved in FAHFA metabolism contribute to sex differences in FAHFA levels. Given the differences in TG estolides, the FAHFA cycling via Atgl, Hsl, and Dgat1 might be affected by sex hormones (15-18).

Emont et al. published a single-cell atlas of mouse white adipose tissue, including data for both sexes (35). Visualization of FAHFA-related genes showed that *Adtrp* (and the other acylglycerol metabolism-related genes) is expressed only in adipocytes and not in other cell types within WAT (Figure 6A). However, the other FAHFA hydrolase *Aig1* is expressed mainly in epithelial and endothelial cells. The same panel clustered on sex showed that the overall expression of *Adtrp* and *Aig1* is comparable among the sexes and that females have higher expression of acylglycerol metabolism-related genes (Figure 6B). Clustering on WAT depot location showed different distributions of gene expression in inguinal WAT (subcutaneous), periovarian, and perigonadal (male) WAT (Figure 6C) (35). Analysis of the cellular composition of WAT suggests that sex differences in FAHFA levels could be influenced by many players within the local environment of WAT, favoring synthesis and disfavoring degradation in females.

**Figure 6.**
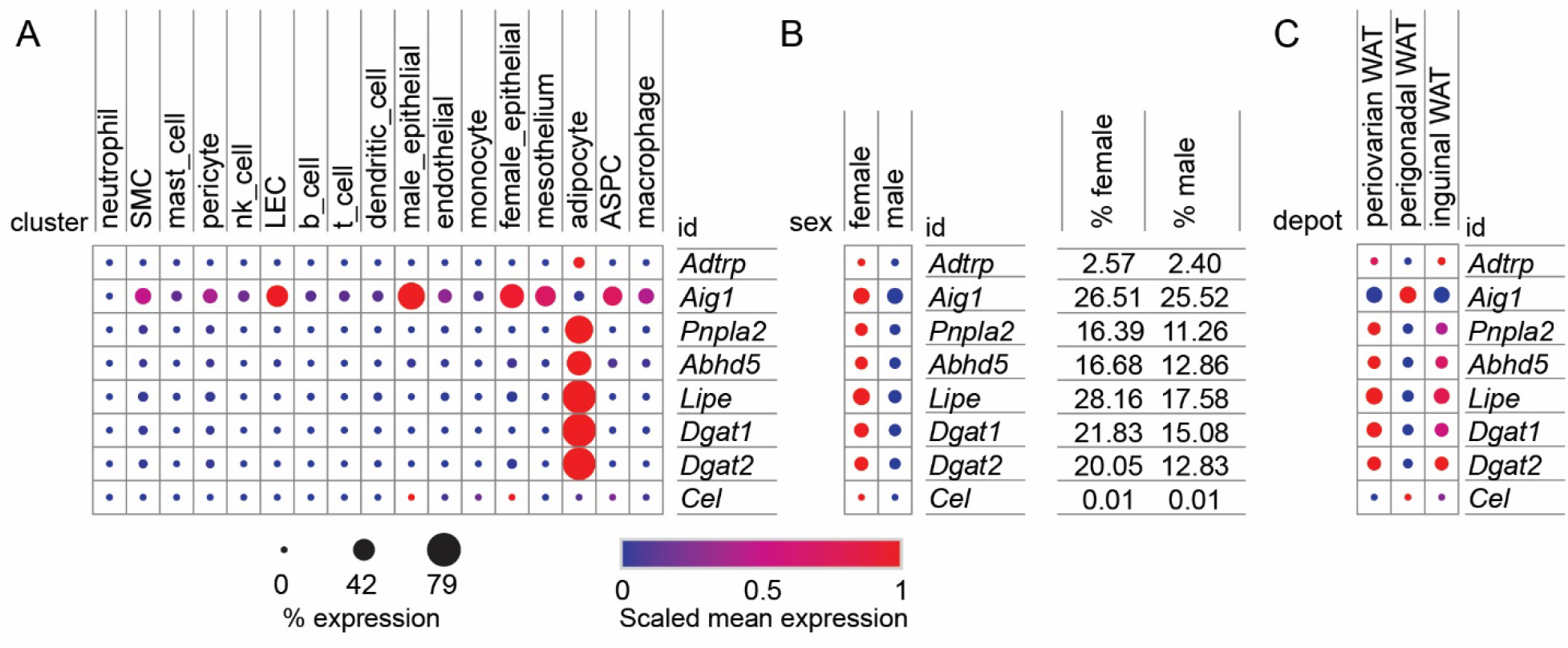
Visualization of FAHFA-related genes in WAT using single-cell mouse WAT atlas (35). A: Dot plot of selected mouse genes within the cluster of cell types. Scaling is relative to each gene’s expression across all cells in a given annotation selection. *Abhd5*, Alpha/beta Hydrolase Domain Containing 5, CGI-58 Atgl co-factor; *Dgat1*, Diacylglycerol O-Acyltransferase 1; *Dgat2*, Diacylglycerol O-Acyltransferase 2; *Lipe*, Hormone-Sensitive Lipase; *Pnpla2*, Patatin Like Phospholipase Domain Containing 2, Atgl; ASPC, adipose stem and progenitor cells; LEC, lymphatic endothelial cells; SMC, vascular smooth muscle cells; B: The same data clustered according to sex. The percentage of gene expression is provided to highlight the difference between males and females. C: The same data clustered according to WAT depot location.

### Perspectives and Significance

Sex-dependent levels of FAHFAs in specific WAT depots could locally modulate adipocyte sensitivity to insulin. This regulation could be altered in obesity by impaired steroidogenesis in adipose tissue (40). Nelson et al. published a preprint showing sex-based heterogeneity of FAHFAs in trained runners (42). Circulating FAHFAs were increased in females in a manner modulated by specific adipose depot sizes, blood glucose, and lean body mass (42). Another preprint by Yan et al. (43) shows that Aig1 is involved in obesity-induced inflammation and insulin resistance in WAT. Future studies using human cohorts and different fat depots will be needed to explore sex-specific FAHFA metabolism in the WAT cellular microenvironment and its impact on systemic metabolism.

## ASSOCIATED CONTENT

Supplementary figures and tables: Figure S1: TG estolides with full names. Table S1, Table S2 – data for volcano plots in Figure 3. Lipidomics Minimal Reporting Checklist.

## DECLARATIONS

### Ethics approval and consent to participate

All animal procedures were approved by Institutional Animal Care and Use Committees of the Oklahoma Medical Research Foundation and by the Institutional Animal Care and Use Committee (IACUC) and the Committee for Animal Protection of the Ministry of Agriculture of the Czech Republic (Approval Number: 81/2016) for the care and use of laboratory animals.

### Availability of data and materials

The datasets during and/or analysed during the current study are available from the corresponding author upon reasonable request.

### Authors’ contributions - CRediT

Conceptualization: F.L. and O.K.; Data curation: V.D. and M.Ro.; Formal analysis: M.R., V.D., M.P., M.V. and O.K.; Funding acquisition: M.Ro., F.L. and O.K.; Investigation: M.R., C.L., M.P. and O.K.; Methodology: M.R., V.D., M.P., M.V. and O.K.; Project administration: M.Ro., A.S., F.L. and O.K.; Resources: C.L., M.Ro., A.S., F.L. and O.K.; Software: M.V. and O.K.; Supervision: O.K.; Validation: M.Ro. and F.L.; Visualization: M.R., M.V. and O.K.; Writing – original draft: M.R. and O.K.; Writing - review & editing: M.R., V.D., C.L., M.P., M.V., M.Ro., A.S., F.L. and O.K.;

## Acknowledgments

The computational resources were supplied by the project “e-Infrastruktura CZ” (e-INFRA LM2018140) provided within the program Projects of Large Research, Development, and Innovations Infrastructures. The data were acquired at the Metabolomics Core Facility at the Institute of Physiology of the Czech Academy of Sciences. Project NU21-01-00469 provided mouse samples. The authors acknowledge Charles University for supporting PhD student M.R.

